# HiCSR: a Hi-C super-resolution framework for producing highly realistic contact maps

**DOI:** 10.1101/2020.02.24.961714

**Authors:** Michael C. Dimmick, Leo J. Lee, Brendan J. Frey

## Abstract

**Motivation:** Hi-C data has enabled the genome-wide study of chromatin folding and architecture, and has led to important discoveries in the structure and function of chromatin conformation. Here, high resolution data plays a particularly important role as many chromatin substructures such as Topologically Associating Domains (TADs) and chromatin loops cannot be adequately studied with low resolution contact maps. However, the high sequencing costs associated with the generation of high resolution Hi-C data has become an experimental barrier. Data driven machine learning models, which allow low resolution Hi-C data to be computationally enhanced, offer a promising avenue to address this challenge.

**Results:** By carefully examining the properties of Hi-C maps and integrating various recent advances in deep learning, we developed a Hi-C Super-Resolution (HiCSR) framework capable of accurately recovering the fine details, textures, and substructures found in high resolution contact maps. This was achieved using a novel loss function tailored to the Hi-C enhancement problem which optimizes for an adversarial loss from a Generative Adversarial Network (GAN), a feature reconstruction loss derived from the latent representation of a denoising autoencoder, and a pixel-wise loss. Not only can the resulting framework generate enhanced Hi-C maps more visually similar to the original high resolution maps, it also excels on a suite of reproducibility metrics produced by members of the ENCODE Consortium compared to existing approaches, including HiCPlus, HiCNN, hicGAN and DeepHiC. Finally, we demonstrate that HiCSR is capable of enhancing Hi-C data across sequencing depth, cell types, and species, recovering biologically significant contact domain boundaries.

**Availability:** We make our implementation available for download at: https://github.com/PSI-Lab/HiCSR

**Supplementary information:** Available Online

## 1 Introduction

In recent years, high-throughput chromosome conformation capture (Hi-C) (Lieberman-Aiden *et al*., 2009) has increasingly enabled studies of the three-dimensional (3D) architecture of the genome. Using proximity-based ligation combined with high-throughput sequencing, the Hi-C method produces a genome-wide heat map contact matrix where each value represents the interaction frequency between two loci. The analysis of Hi-C contact matrices has led to significant discoveries on the nature of chromatin substructures such as A/B Compartments (Lieberman-Aiden *et al*., 2009), Topologically Associating Domains (TADs) (Dixon *et al*., 2012), and chromatin loops (Rao *et al*., 2014). Beyond the discovery of architectural substructures, chromatin conformation has been shown to play a significant role in gene regulation and expression (Franke *et al*., 2016; Lupiáñez *et al*., 2015), illustrating the important relationship between genome architecture and cellular functions.

The resolution of a Hi-C matrix is determined by the chosen genomic bin size, with a smaller bin size resulting in a higher resolution. This choice of bin size is typically determined by sequence depth, as an insufficient number of sequence reads for a given resolution results in sparse and noisy Hi-C data. In general, a linear increase in resolution requires a quadratic increase in sequencing depth (Schmitt *et al*., 2016), making high resolution Hi-C data costly to obtain. While Hi-C matrices with high resolutions (≤ 10 Kb) have been generated, the large cost incurred for increasing sequence depth results in an abundance of low resolution datasets (e.g. 40 Kb - 1 Mb). These low resolution datasets encumber the analysis of finer substructures in the 3D genome, as the details of certain substructures such as chromatin loops cannot be accurately identified in low resolution contact maps (Rao *et al*., 2014). There is therefore both a clear advantage and high demand for high resolution Hi-C datasets.

The large amounts of publicly available Hi-C data provides an opportunity for researchers to develop new predictive tools which can both accelerate experimentation and enable new discoveries. In the context of Hi-C, deep learning methods can leverage this large volume of data to computationally increase the resolution of a Hi-C contact matrix in scenarios where the sequence depth is low. This is done by learning a mapping between high and low resolution contact maps, similar to natural image based methods for enhancement and denoising (Ledig *et al*., 2017; Zhang *et al*., 2017). These techniques provide researchers with a means to generate high resolution Hi-C datasets with significantly fewer sequencing reads than would otherwise be required for a given resolution. Although different deep learning based Hi-C enhancement methods have been developed, they fall short in several aspects. Methods which optimized for a Mean Squared Error (MSE), such as HiCPlus (Zhang *et al*., 2018) and HiCNN (Liu T. *et al*., 2019), suffer from a lack of high frequency information resulting in a blurred output. This is caused by an objective function which prefers solutions that are the pixel-wise average of many possible solutions that lie on the plausible image manifold (Mathieu *et al*., 2016). To avoid blurred predictions, hicGAN (Liu Q. *et al*., 2019) and DeepHiC (Hong *et al*., 2019) were proposed. First, hicGAN replaced pixel-wise loss functions with a purely adversarial loss. However, this caused hicGAN predictions to miss details found in the true high resolution data. DeepHiC combined an adversarial loss, pixel-wise loss, and a perceptual loss derived from a VGG-16 loss network (Simonyan and Zisserman, 2015) trained on ImageNet. However, the introduction of this perceptual loss caused unwanted natural image textures in DeepHiC’s predictions not otherwise found in real Hi-C data.

Improving upon these methods, we proposed a novel Hi-C Super-Resolution (HiCSR) framework capable of inferring high resolution Hi-C data from low resolution Hi-C data with high accuracy. This was achieved using a new loss function tailored to the Hi-C enhancement problem. HiCSR optimizes a weighted combination of adversarial loss, pixel-wise L1 loss, and a feature reconstruction loss obtained from the latent representation of a task specific denoising autoencoder (Vincent *et al*., 2008). While previous enhancement methods have failed to address the unique properties of Hi-C data in their evaluations, relying primarily on correlative and image-based metrics, we opt for Hi-C specific metrics of reproducibility as a more meaningful measure of model performance. To our knowledge, this is the first effort to compare a suite of Hi-C super resolution methods in this way. We demonstrate that HiCSR enhanced Hi-C data consistently achieves strong reproducibility scores across cell types and species with respect to the true high resolution Hi-C data, and outperformed all previously proposed super-resolution models in terms of reproducibility. We also find that HiCSR enhanced data is capable of recovering Hi-C specific attributes, such as insulation score values and TAD boundaries in multiple cell types and across species.

## 2 Methods

### 2.1 Unsupervised representation learning of Hi-C data

To begin, we describe an unsupervised approach to learning a representation of high resolution Hi-C data. This approach is motivated by the notion that a good representation of the input data should be robust to noise corruptions. Additionally, a strong performance in this denoising task requires a good feature representation that adequately captures the structure of the input (Vincent *et al*., 2010). We therefore propose to use a denoising autoencoder to learn the Hi-C representation in an unsupervised fashion. We train a model to denoise high resolution Hi-C data and in the process, learn a useful feature representation in the model’s latent space. Specifically, high resolution Hi-C matrix *I^HR^* is first corrupted with zero mean Gaussian noise, producing a noisy version of the input data 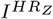:

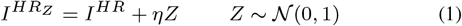

where *η* is a noise corruption factor. The noise corrupted input is then passed through the denoising autoencoder network *ϕ* to produce the reconstruction 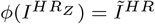. An overview of this setup is illustrated in Fig. 1A. For each new Hi-C sample provided to the network during training, a new noise sample is produced from the noise generating distribution and added to the input. These noise corruptions are only added during the training phase of the denoising autoencoder. Once the model is trained, representations are extracted from the noise-free data. The denoising autoencoder is trained to reconstruct the noise-free *n* × *n* input by minimizing the MSE between the original high resolution contact matrix and the predicted reconstruction:

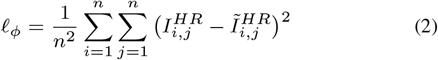

**Fig. 1.**
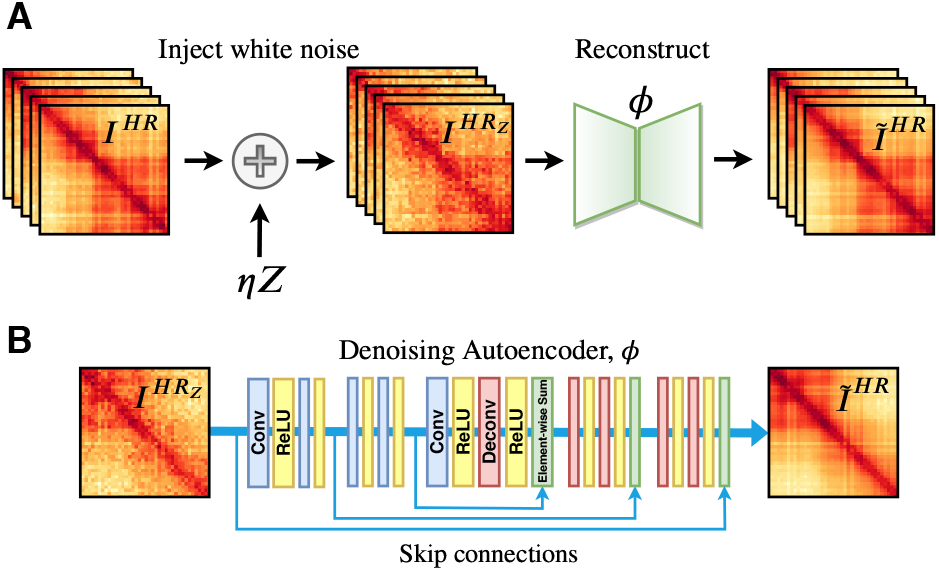
Overview of Hi-C representation learning. (A) Denoising autoencoder setup. High resolution Hi-C data is first corrupted by adding zero mean Gaussian noise to the input. The corrupted data is then passed into the reconstruction network *ϕ*. The network *ϕ* learns to denoise corrupted Hi-C data and predict a noise-free reconstruction. (B) Architecture of the denoising autoencoder. The model consists of five convolutional layers followed by five deconvolutional layers with skip connections every other layer. The network takes noisy high resolution Hi-C data as input and infers a noise-free reconstruction.

The denoising autoencoder employs a slightly modified image restoration architecture from Xiao-Jiao *et al*., 2016 shown in Fig. 1B. The encoder and decoder portions are each made up of five convolutional layers with ReLU (Nair and Hinton, 2010) activation and a tanh output function. He initialization (He *et al*., 2015) is used for all convolutional and deconvolutional layers. Throughout the model, every other layer also includes a symmetric skip connection which performs an element-wise sum between the encoder’s post-activation convolutional output with its counterpart in the decoder. These skip connections aid in the flow of gradients during backpropagation, as well as enable the model to pass fine details of the Hi-C matrix to the decoding layers allowing for the end-to-end learning of deeper networks to be more effective and efficient (He *et al*., 2016). Each convolutional and deconvolutional layer consists of 64 filters of size 3 × 3. Once trained, the denoising autoencoder is able to compute a feature representation of Hi-C data of any input size, as the model is fully convolutional.

### 2.2 HiCSR framework

In this section, we provide a detailed description of the HiCSR framework. HiCSR is a Hi-C enhancement model which combines a Generative Adversarial Network (GAN) (Goodfellow *et al*., 2014) architecture with a denoising autoencoder loss network to predict accurate high resolution Hi-C data from insufficiently sequenced samples. Given pairs of low and high resolution Hi-C contact matrices (*I^LR^, I^HR^*), where *I^LR^* is a down-sampled version of *I^HR^* (e.g. 16× fewer sequence reads), HiCSR produces a prediction *I^SR^* of the high resolution sample. An overview of the framework is shown in Fig. 2A and additional information on model parameters can be found in Supplementary Table 1.

**Fig. 2.**
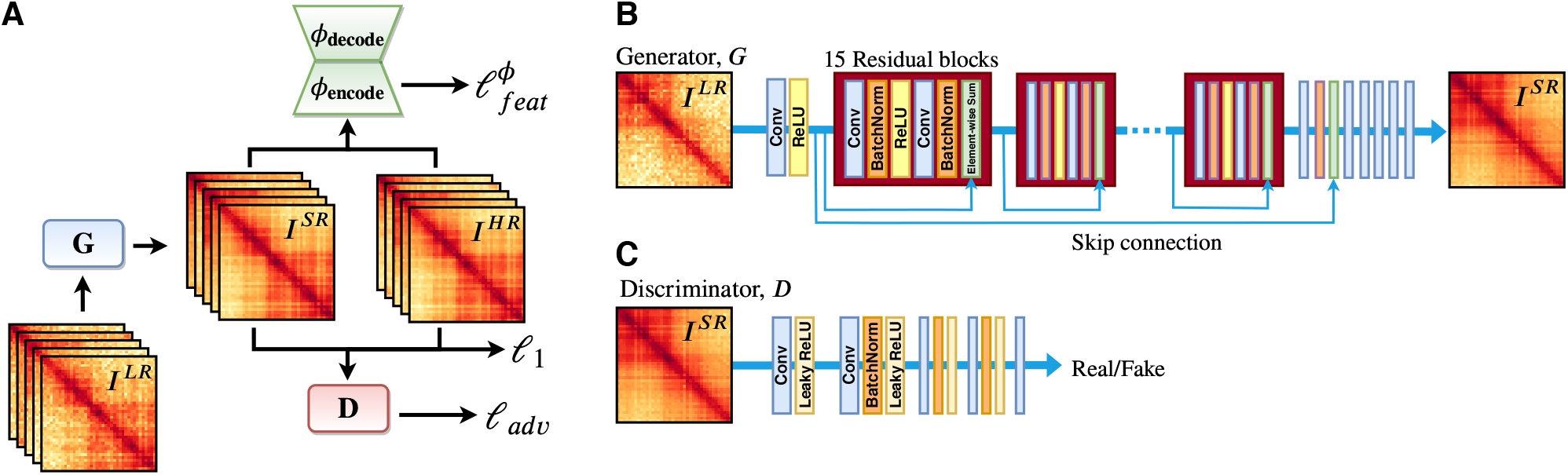
Overview of the HiCSR framework. (A) HiCSR employs an adversarial training strategy, augmented by two additional loss functions. First, low resolution Hi-C data is passed to the generator network *G* which produces a super-resolved output. Both the super-resolved output and true high resolution Hi-C data are then used as inputs to compute the three components of the generator’s loss function. The discriminator network *D* predicts whether each input sample comes from either the true high resolution Hi-C data distribution or from the generator, and its performance on this task is used to compute an adversarial loss *ℓ_adv_*. Additionally, a denoising autoencoder loss network *ϕ* then computes a feature reconstruction loss 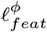 between the inputs. Finally, the inputs are used to compute the pixel-wise L1 loss *ℓ*_1_. (B) Generator network *G* consists of 15 residual blocks with a skip connection. The generator enhances low resolution Hi-C data producing the super-resolved output. (C) Discriminator network *D* is fully convolutional and predicts the probability that the sample came from real high resolution Hi-C data, as opposed to super-resolved Hi-C data produced by the generator.

HiCSR optimizes for a weighted combination of losses: an adversarial loss produced from the discriminator of a GAN, a feature reconstruction loss computed from the feature representations of a denoising autoencoder, and a pixel-wise L1 loss. The total objective function of the generator is given by:

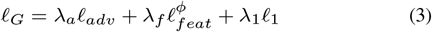

where *λ_a_*, *λ_f_*, and *λ*_1_ are scaling constants. Each component of the loss function focuses on a specific and desirable aspect of enhancement. The adversarial loss *ℓ_adv_* ensures that the generator favours outputs which lie on the true high resolution Hi-C data manifold, encouraging visually convincing solutions. The feature reconstruction loss 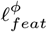 ensures that HiCSR enhanced Hi-C data shares accurate feature representations with true high resolution Hi-C data. Finally, the L1 loss *ℓ*_1_ encourages similarity on the level of individual pixels. The minimization of the combined losses ensures that HiCSR generates enhanced Hi-C data that is both accurate and visually convincing.

#### 2.2.1 Adversarial loss

The GAN training method employs two neural networks to produce synthetic samples which appear to come from the desired data generating distribution. The first network called the generator, takes samples from an input distribution, and through a series of non linear transformations produces synthetic samples which appear to have been drawn from the desired sample distribution. The second network is the discriminator, which takes samples from either the generator or the training set as input, and attempts to classify them as having either come from the generator or the training set. The two networks are trained in an alternating fashion and play an adversarial game. As the generator improves in its ability to create realistic samples, the discriminator improves its ability to distinguish between the training distribution and the generator’s output. Training is considered successful when the generator has learned to create synthetic samples which the discriminator cannot accurately distinguish from real samples of the desired distribution.

Forming this description mathematically in the context of Hi-C enhancement, we define the generator network *G* (Fig. 2B) and discriminator network *D* (Fig. 2C), parameterized by *θ_G_* and *θ_D_* respectively. During training, the generator network takes low resolution Hi-C data *I^LR^* as input and produces super-resolved Hi-C data *I^SR^* = *G*(*I^LR^*) as output. The discriminator network takes both the super-resolved matrix *I^SR^* produced from the generator and true high resolution Hi-C data *I^HR^* as input, and classifies the input sample as either real (high resolution) or fake (a sample produced by the generator). The objective function of the two networks can then be written as a minimax game:

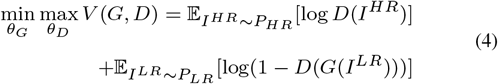

where *P_HR_* and *P_LR_* are the data generating distributions of high and low resolution Hi-C data, respectively. The discriminator’s classification performance on this task is used to produce the first component of the generator’s loss function, the adversarial loss. We used the standard generator loss (Goodfellow *et al*., 2014) over all *n* = 1,…, *N* training samples:

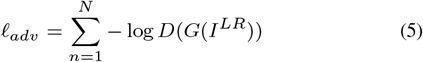

The generator uses a fully convolutional architecture so that once trained, the model allows for any size of Hi-C data to be enhanced. The generator implements residual learning (He *et al*., 2016) and consists of 15 residual blocks as well as a skip connection. Each convolutional layer within the residual blocks consists of 64 (3 × 3) filter maps. The discriminator model is also fully convolutional and uses Leaky ReLU activations. Both models leverage Batch Normalization (Ioffe and Szegedy, 2015) for regularization and to reduce training time.

#### 2.2.2 Feature reconstruction loss

Inspired by previous work outlining the use of loss networks (Johnson *et al*., 2016; Gatys *et al*., 2015), the HiCSR framework employs a denoising autoencoder (described in Sec. 2.1) loss network *ϕ* pretrained to reconstruct noise corrupted high resolution Hi-C data. To do so, both the enhanced Hi-C matrix *I^SR^* and the true high resolution Hi-C matrix *I^HR^* are individually passed through the loss network, and their reconstructions: 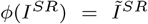, and 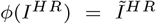 are computed. Once both predictions are made, the intermediate feature representations from all encoder layers are extracted.

Using these intermediate feature representations, we then computed a feature reconstruction loss 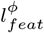 which measures the similarity between the feature maps obtained from passing the super-resolved Hi-C data and true high resolution Hi-C data through the denoising autoencoder. Specifically, the feature reconstruction loss is the sum of the squared normalized Euclidean distances between the pre-activation feature representations of the true high resolution matrix *I^HR^*, and enhanced matrix *G*(*I^LR^*) across all *K* layers of the encoder network:

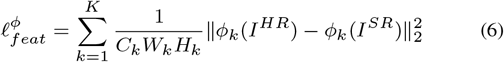

where *ϕ_k_* is the pre-activation feature representation of the denoising autoencoder at layer *k* of the encoder. For each sample, this similarity is computed for feature maps of shape *C_k_* × *W_k_* × *H_k_* for each of the *k* = 1,…,*K* encoder layers between two separate denoising autoencoder inputs.

#### 2.2.3 Pixel-wise L1 loss

While both the adversarial and feature reconstruction losses discussed thus far encourage useful properties for Hi-C enhancement, they have no explicit criteria that encourages a faithful prediction in the pixel-wise sense. For this reason, we include a Mean Absolute Error (MAE) / L1 loss computed between the super-resolved and true Hi-C high resolution Hi-C data:

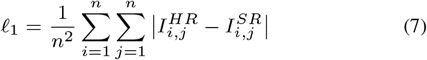

### 2.3 HiCSR dataset generation and preprocessing

The dataset used to train and evaluate HiCSR requires pairs of high and low resolution Hi-C data. To obtain these sets of matrices, we began with a database of sequence reads for a given cell type and generated the low resolution data through a uniform random down-sampling of the original aligned reads by a factor of 16. Both the original (high resolution) and down-sampled reads were then processed into low and high resolution contact maps using the Hi-C processing pipeline, HiC-Pro (Servant *et al*., 2015) with default settings. As there are few meaningful interchromosomal interactions, only the intrachromosomal contact matrices are considered.

Both sets were then normalized by sequence depth to remove model dependency on the total number of raw interactions. We define the matrix *M^c^* as the raw contact matrix of chromosome *c*, and performed a log transform on the contact matrices given by:

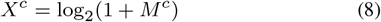

We then applied a linear transform 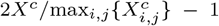, normalizing the matrices to the range [−1, 1] for each chromosome. The normalized contact matrices were then cropped to generate low resolution *n*×*n* sub-matrices and their corresponding high resolution counterparts for each chromosome. As most meaningful interactions occur within TADs, and the majority of TADs are less than 1 Mb in size within the human genome, interactions with a genomic distance greater than 2 Mb (far from the matrix diagonal) were discarded.

### 2.4 HiCSR evaluation

Previously, all Hi-C enhancement models have been evaluated and compared using image and correlation based measures. With respect to image based approaches, MSE tends to favour blurred solutions (Mathieu *et al*., 2016), and the applicability of the Structural Similarity (SSIM) index (Wang *et al*., 2004) to Hi-C similarity is questionable, as it was designed for evaluating the perceptual quality of natural images. With respect to correlation measures, it has been shown that two unrelated biological samples can have a high Pearson correlation coefficient and furthermore, it is possible to find higher Pearson and Spearman correlations between unrelated samples than those between true biological replicates (Yang *et al*., 2017; Yardimci *et al*., 2017). In these cases, both Pearson and Spearman Correlation metrics fail to account for the unique spatial structure found in Hi-C data, as well as the “distance effect” (propensity for increased contact frequency between loci at small linear distances along the genome (Lieberman-Aiden *et al*., 2009)) seen in all Hi-C datasets.

We therefore focus on a more biologically significant measure, evaluating HiCSR on Hi-C specific metrics which quantify the reproducibility of Hi-C samples. We employed a Hi-C reproducibility software package (Yardimci *et al*., 2017) which combines four different software tools developed by members of the ENCODE Consortium to compute reproducibility/similarity scores: GenomeDISCO (Ursu *et al*., 2018), Hi-C Spector (Yan *et al*., 2017), HiCRep (Yang *et al*., 2017), and QuASAR-Rep (Sauria and Taylor, 2017). Each of these methods propose a unique perspective for scoring the similarity between two contact maps. GenomeDISCO performs a series of random walks on a network created from the Hi-C data in order to first smooth the contact map. Similarity is then computed as the difference between the smoothed contact maps. Hi-C Spector focuses on spectral reproducibility, comparing the eigen-decomposition of the Laplacian matrix between samples using a weighted difference of eigenvectors. The HiCRep method accounts for both contact map sparsity and the “distance effect” in its measure. HiCRep smooths the contact map using a 2D mean filter and then stratifies the contact map by genomic distance, computing similarity as the strata-weighted correlation between contact maps. QuASAR-Rep computes similarity by testing for the assumption that spatially proximal loci will produce similar contact frequencies throughout the genome.

These measures for reproducibility provide a tool for computing contact map similarity which is designed with the unique aspects of Hi-C data in mind. These software tools allow researchers to quantitatively validate the quality and reproducibility between a previously validated sample and a new experimental sample to confirm high quality experimental results. In the context of Hi-C enhancement, Hi-C reproducibility measures provide a logical basis for the comparison between a super-resolved matrix and the true high resolution Hi-C data.

We compared HiCSR’s performance on each of the four described reproducibility metrics against all previously proposed Hi-C enhancement methods including HiCPlus, HiCNN, hicGAN, and DeepHiC. On this evaluation, a successful Hi-C super-resolution model would therefore score a high reproducibility value across all four measurements. In the same way, we also evaluated HiCSR’s ability to transfer information learned on a training set cell type to new cell types. We retrained HiCSR on a new data set from a different cell type, and evaluated the reproducibility scores of the enhanced Hi-C data on cell types that were previously unseen.

Despite the aforementioned weaknesses of image and correlation based evaluations of Hi-C data, for completeness we also compared HiCSR to other Hi-C enhancement methods using MSE, MAE, Peak Signal-to-noise Ratio (PSNR), as well as both Pearson and Spearman correlation. For this evaluation, each metric is computed as a function of genomic distance and the results are averaged over distances less than 2.0 Mb apart.

## 3 Results

### 3.1 Denoising autoencoder produces high fidelity reconstructions of Hi-C data

Before evaluating the enhancement capability of the HiCSR model, we confirmed that the trained denoising autoencoder loss network (used to compute a feature reconstruction loss) is able to accurately reconstruct high resolution Hi-C data. The denoising autoencoder was trained on 10 Kb resolution Hi-C data from the GM12878 cell type using all available paired-end Hi-C reads downloaded from the Gene Expression Omnibus (GEO) database (accession GSE63525) (Rao *et al*., 2014). The Denoising autoencoder network uses chromosomes 1-16 for training, 17 and 18 for hyper-parameter tuning, and 19-22 for evaluation. The model was trained on a total of 70484 overlapping sub-matrices of size *n* × *n*, where *n* = 40 (i.e. Hi-C patches of size 0.4 × 0.4 Mb). The denoising autoencoder was trained over 600 epochs using the Adam optimizer (Kingma *et al*., 2014) with a batch size of 256, a learning rate of 5×10^−3^, and a noise corruption factor of *η* = 0.1.

We evaluated the denoising autoencoder’s reconstruction capability from several perspectives. When inspecting the denoising autoencoder’s performance qualitatively (Fig. 3A), we found that the reconstruction captures many of the visual intricacies of the original Hi-C data, and produces a high fidelity reconstruction. We then compared samples from the high resolution Hi-C data to the reconstructed samples of Hi-C data produced from the denoising autoencoder using MSE (Fig. 3B). To further validate, we grouped all test chromosomes together, binned the MSE values according to short-range interactions (0.00-0.25 Mb), mid-range interactions (0.25-1.00 Mb) and long-range interactions (1.00-2.00 Mb), taking the average within bins. For short-, mid-, and long-range interactions, the denoising autoencoder achieved a MSE of 0.013, 0.025, and 0.033, respectively. We found that the reconstructed matrix samples achieved low error across all four test chromosomes, and that the model produced particularly small errors at low genomic distances. This is a desirable outcome for Hi-C matrix reconstructions as the most significant and least noisy interactions occur at low genomic distances.

**Fig. 3.**
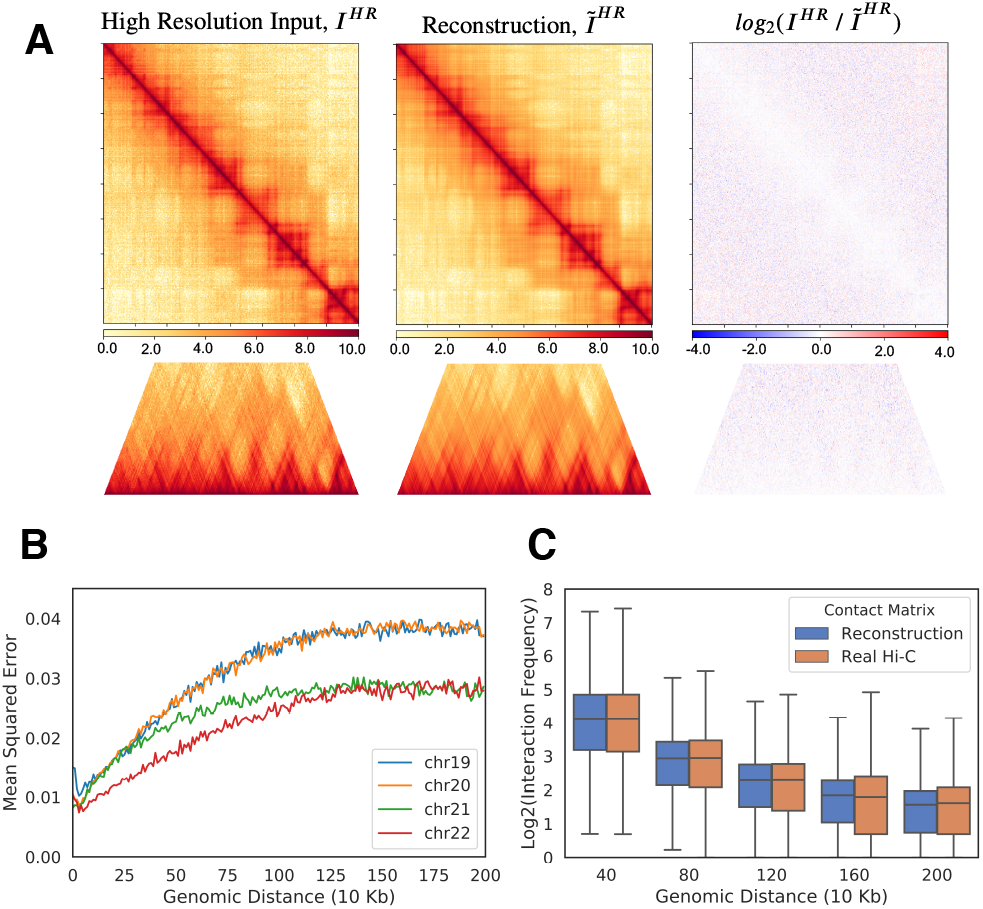
Evaluation of denoising autoencoder reconstructions in the GM12878 cell type. (A) Sample denoising autoencoder prediction from chromosome 20 (2.0 Mb - 4.0 Mb). Visualizations depict the original high resolution Hi-C data (left), Hi-C data reconstruction (middle) and the log ratio between them (right). (B) Mean Squared Error between the true Hi-C data and the reconstruction as a function of genomic distance for chromosomes 19-22. The reconstruction from the denoising autoencoder achieves consistently low error on all four test chromosomes. (C) Comparison of interaction frequencies between the denoising autoencoder reconstruction and real Hi-C data, binned by genomic distance. The denoising autoencoder-produced reconstructions achieve similar interaction frequencies to that of real Hi-C data across a range of genomic distances.

For additional comparison we also evaluated the contact frequency profiles of both real Hi-C data and reconstructed Hi-C data from the denoising autoencoder. We computed the base 2 logarithm of the raw interaction frequencies and grouped them into 0.4 Mb sized bins. We then compared the distributions of the resultant binned interaction frequencies between real Hi-C data from the test chromosomes and their corresponding reconstructions (Fig. 3C). We found that both the real and reconstructed Hi-C data have similar interaction frequency profiles at a variety of genomic distances, and the reconstructed matrix correctly matches the exponential drop off in contact frequency seen in real Hi-C data.

### 3.2 Denoising autoencoder learns a meaningful representation of high resolution Hi-C data

Next, we aimed to evaluate the learned representations of the denoising autoencoder. In line with the proposed feature reconstruction loss used in the HiCSR framework, we analyzed the feature space of the model using the same setup used to evaluate the denoising autoencoder reconstructions in the previous section. To determine if the learned feature representations are useful for Hi-C enhancement tasks, we inspected the pre-activation feature representations at each layer of the encoder when high resolution Hi-C data is passed as input. For select filters within the model, we discovered that the corresponding feature maps captured key high frequency textures which are ubiquitous in Hi-C contact maps (Fig. 4A). Specifically, we found filters which preserved the speckled textures typically found in Hi-C data, as well as activation maps which emphasize high frequency hatching patterns in both the vertical and horizontal directions. These learned representations encapsulate some of the lower-order structures found within Hi-C contact matrices, and contribute to the overall visual particularities of the data. The reconstruction of these textures is a desirable characteristic of a Hi-C super-resolution model, as they allow the super-resolved outputs to faithfully preserve the high frequency information that is captured in real Hi-C data. It is therefore a good indication that the denoising autoencoder has learned useful feature representations for Hi-C enhancement.

**Fig. 4.**
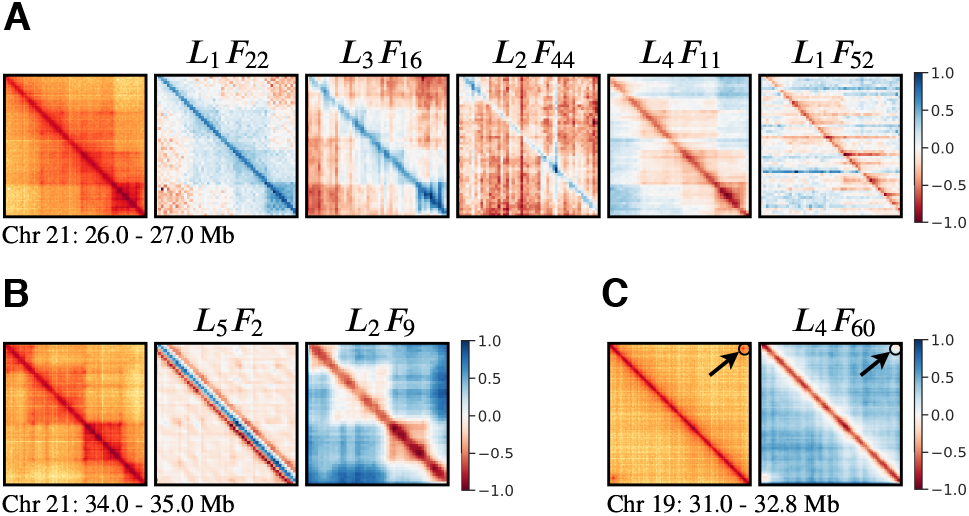
Visualization of the denoising autoencoder feature representations in the GM12878 cell type depicting High resolution Hi-C data input and select examples of the accompanying pre-activation feature maps. (A) Sample from chromosome 21 (26.0 - 27.0 Mb). Select pre-activation feature maps *F* from the encoder layers *L* capture the high frequency content and textures found in Hi-C data. Sample feature maps illustrate high frequency speckling textures (*L*_1_*F*_22_), vertical hatching patterns (*L*_3_*F*_16_, *L*_2_*F*_44_) and horizontal hatching patterns (*L*_4_*F*_11_, *L*_1_*F*_52_). (B) Sample from chromosome 21 (34.0 - 35.0 Mb). Pre-activation maps of the encoder exhibit interpretable substructures found in Hi-C data. Feature maps emphasize self-interacting loci (*L*_5_*F*_2_) and distinguish Topologically Associating Domains (TADs) from other interactions (*L*_2_*F*_9_). (C) Samples from chromosome 19 (31.0 - 32.8 Mb). Pre-activation maps of the encoder contains representations of chromatin loops (*L*_4_*F*_60_).

Beyond simple texture information, we discovered that the denoising autoencoder was also able to capture higher-order information. We discovered that the denoising autoencoder feature representations captured interpretable chromatin substructures known to exist in true high resolution Hi-C data. Select filters from the encoder layers were able to capture structural information found in high resolution Hi-C data, and the representations emphasize biologically interpretable features such as self-interacting loci, and TADs (Fig. 4B). We find that the denoising autoencoder is also capable of representing the relative spike in interaction frequencies corresponding to chromatin loops (Fig. 4C). The preservation of these substructures within the feature representations are a promising sign that the denoising autoencoder has learned a meaningful representation of high resolution Hi-C data, and would therefore provide benefit to the Hi-C enhancement process.

### 3.3 HiCSR-enhanced data consistently achieves high reproducibility scores across cell types

HiCSR was trained on pairs of low and high resolution Hi-C data from the GM12878 cell type with the same dataset split used for the denoising autoencoder (1-16 for training, 17 and 18 for validation). In total, HiCSR is trained with 70484 pairs of sub-matrices of size *n × n*, where *n* = 40. The generator and discriminator training was done in an alternating fashion over 500 epochs using the Adam optimizer with a batch size of 128, and a learning rate of 10^−5^. The HiCSR training procedure took 48 hours on a single NVIDIA Tesla P100 GPU. The LeakyReLU activation used in the discriminator was implemented with *α* = 0.2. Scaling factors *λ_a_*, *λ_f_*, and *λ*_1_ were chosen through cross-validation as 2.5 × 10^−3^, 1.0, and 1.0, respectively. Once trained, the discriminator and denoising autoencoder were discarded, and Hi-C super-resolution predictions were made with the generator alone.

We first evaluated HiCSR on image and correlation based metrics in a similar fashion to previous methods, computing the MSE, Peak Signal-to-noise Ratio, MAE, Pearson, and Spearman Correlation averaged over genomic distances ≤ 2.0 Mb (Table 1). We evaluated HiCSR against a suite of previously developed deep learning based Hi-C enhancement methods, including HiCPlus, HiCNN, hicGAN and DeepHiC. For DeepHiC and hicGAN, which utilize normalization to remove the model’s dependency on sequence depth, we used the provided relevant pretrained models. As HiCPlus and HiCNN prescribe no normalization and are therefore sensitive to sequence depth, we retrained the models according to the training methods described in the original papers (Zhang *et al*., 2018; Liu T. *et al*., 2019). Although HiCNN performs best on most of these metrics, HiCSR achieved state-of-the-art performance on MAE, and competitive performance overall.

**Table 1.** Image based metrics Mean Squared Error (MSE), Peak Signal-to-noise Ratio (PSNR), and Mean Absolute Error (MAE), as well as Pearson and Spearman Correlation metrics within a genomic distance of 2.0 Mb, averaged over chromosomes 19-22 in the GM12878 cell type. All metrics were computed between true high resolution and enhanced Hi-C data.

| Model | MSE | PSNR (dB) | MAE | Pearson | Spearman |
| --- | --- | --- | --- | --- | --- |
| High resolution | 0.00 | ∞ | 0.00 | 1.00 | 1.00 |
| 16x down-sample | 3.596 | 12.96 | 1.610 | 0.616 | 0.639 |
| HiCPlus | 0.146 | 26.89 | 0.279 | 0.934 | 0.900 |
| HiCNN | **0.118** | **27.81** | 0.245 | **0.943** | **0.909** |
| hicGAN | 0.270 | 24.20 | 0.384 | 0.877 | 0.842 |
| DeepHiC | 0.179 | 26.01 | 0.313 | 0.932 | 0.891 |
| HiCSR (ours) | 0.143 | 26.97 | **0.244** | 0.926 | 0.889 |

We then moved to evaluate HiCSR on a more biologically relevant metric, testing the HiCSR model on four different software tools which measure the intrachromosomal reproducibility between the true high resolution Hi-C data and an enhanced Hi-C matrix. These reproducibility methods: GenomeDISCO, HiC-Spector, HiCRep, and QuaSAR-Rep, assign a score in the range [0, 1] to an enhanced Hi-C matrix indicating the reproduction quality between the input matrix and the true high resolution Hi-C contact map. We computed a reproducibility score for each of the four methods for the six test chromosomes (19-22, X, Y) resulting in a total of 24 separate evaluations for each model (Supplementary Table 2). We found that HiCSR enhanced Hi-C matrices achieved a high score for each of the reproducibility metrics. Averaging over the test chromosomes, HiCSR enhanced data achieved a score of 0.927 from GenomeDISCO, 0.950 from HiC-Spector, 0.973 from HiCRep, and 0.980 from QuASAR-Rep. Across all reproducibility scoring methods, HiCSR enhanced data received consistently high scores, performing the best in 20 out of 24 reproducibility comparisons. We found that HiCSR outperforms all other Hi-C super-resolution models in terms of reproducibility when tested across all four metrics. A comparison of the mean performance on these metrics between Hi-C enhancement models (Fig. 5A) shows that HiCSR (with an average score of 0.958) provides a significant improvement over all other models (*p* < 0.05, one-sided independent t-test), including HiCNN which has the second highest average reproducibility score of 0.931. We also found that when comparing reproducibility scores across the four metrics (Fig. 5B), previous Hi-C super-resolution methods perform poorly in comparison to HiCSR using the GenomeDISCO and HiC-Spector metrics. This indicates that HiCSR is able to better recover the high frequency information found in experimental Hi-C data.

**Fig. 5.**
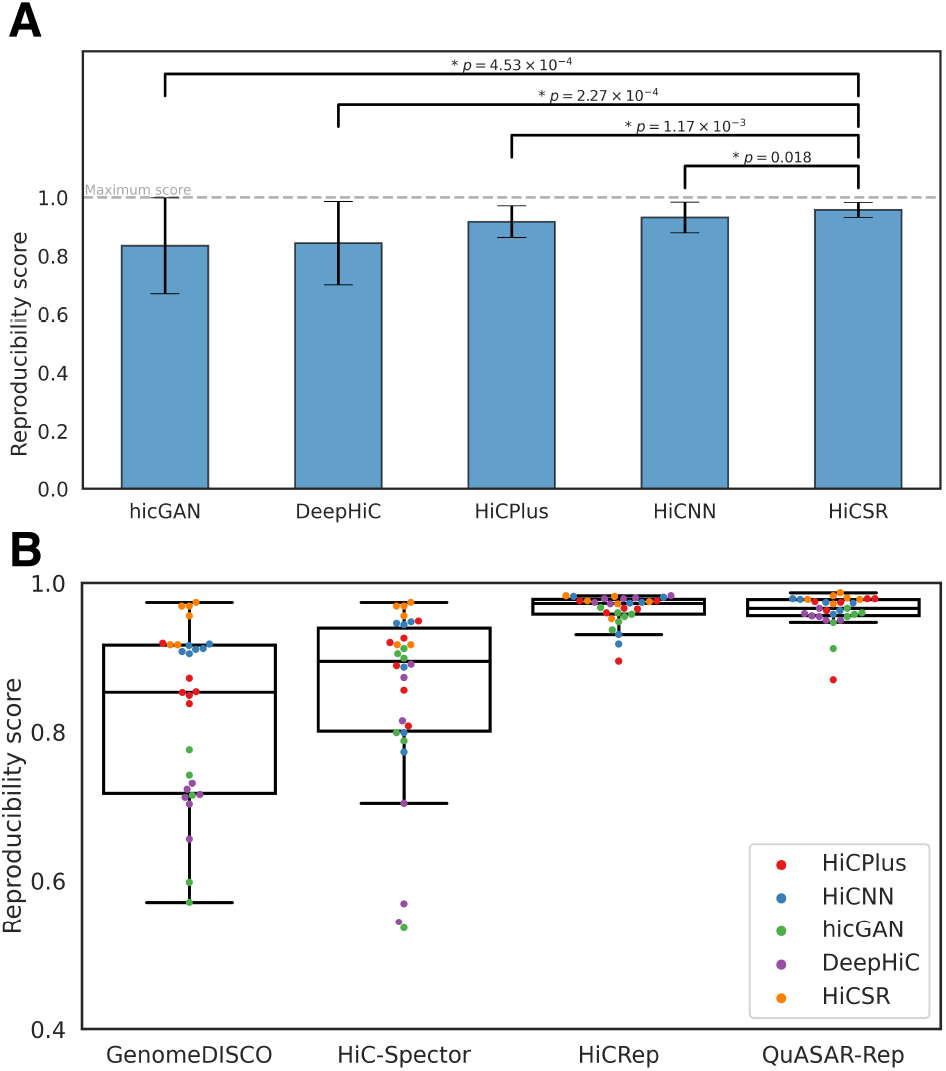
(A) Mean reproducibility scores for all Hi-C enhancement models from left to right: hicGAN = 0.84, DeepHiC = 0.84, HiCPlus = 0.92, HiCNN = 0.93 and HiCSR = 0.96. HiCSR outperforms all other super-resolution models in terms of reproducibility (* One-sided independent t-test). Error bars signify standard deviation. (B) Comparison between super-resolution model performances on each of the four reproducibility metrics, GenomeDISCO, HiC-Spector, HiCRep, and QuASAR-Rep. While previous models perform relatively poorly on GenomeDISCO and HiC-Spector, HiSR performs equally well across all metrics.

We provide a visual comparison of all model enhancements illustrated in Fig. 6A. This comparison highlights the various enhancement artifacts produced by current Hi-C super-resolution methods. We note that HiCPlus- and HiCNN-enhanced matrices tend be overly smooth and blurred compared to real Hi-C data. With hicGAN-enhanced data, the predictions are error prone due to its lack of a pixel-wise loss. Finally, with DeepHiC-enhanced data, we found “wave-like” artifacts produced which do not occur in real Hi-C data. We speculate that these optimization artifacts are caused by the use of a pretrained VGG based perceptual loss, encouraging textures found in the natural image domain on which VGG was trained. In contrast, we found that the HiCSR framework is able to produce highly realistic contact maps which more accurately capture the realistic textures found in experimental Hi-C data. Additional visual samples emphasizing the different optimization artifacts of each model are provided in Supplementary Fig. 1.

**Fig. 6.**
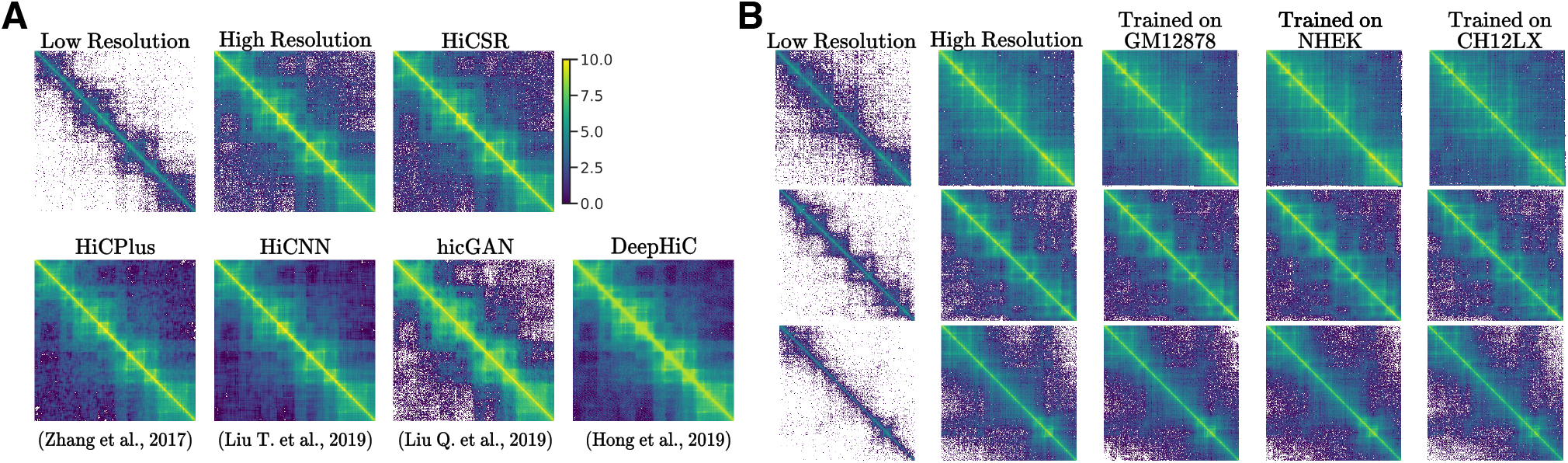
(A) Hi-C super-resolution visual comparisons on a log scale. Samples shown are from chr21: 33.00 - 37.00 Mb at 10 Kb resolution in the GM12878 cell type. Hi-C contact maps depicted are as follows from left to right: 16x down-sampled low resolution, true high resolution, HiCSR (top), HiCPlus, HiCNN, hicGAN, and DeepHiC (bottom). (B) Visual comparison on a log scale between HiCSR predictions when trained on the GM12878, NHEK, and CH12LX cell types. Samples shown are from chr21: 37.00 - 41.00 Mb (top), chr21: 46.00-50.00 Mb (middle), and chr21: 16.00 - 20.00 Mb (bottom) at 10 Kb resolution.

We also tested HiCSR’s ability to transfer super-resolution mappings learned from one cell type to another previously unseen cell type. To do this, we made use of 10 Kb resolution Hi-C data from two additional cell types: NHEK (human) and CH12LX (mouse). For each cell type we used all available paired-end reads downloaded from the Gene Expression Omnibus (GEO) database (accession GSE63525). We again computed reproducibility scores (Table 2) when HiCSR was trained on NHEK or CH12LX, in addition to the original, matching GM12878. Visual samples of HiCSR trained on different cell types but tested on the same GM12878 cell line are provided in Fig. 6B. Although there is a slight performance degradation when training and testing on different cell types, we found that HiCSR is able to produce enhanced Hi-C data that achieves both visual realism and high reproducibility scores across all four measures (Supplementary Fig 2.).

**Table 2.** Reproducibility scores from GenomeDISCO, HiC-Spector, HiCRep, and QuASAR-Rep with HiCSR trained on the GM12878, NHEK and CH12LX cell types, and evaluated on chromosomes 19 - 22 of the GM12878 cell type. All reproducibility scores were computed between true high resolution and enhanced Hi-C data.

| GM12878 | GenomeDISCO | HiC-Spector | HiCRep | QuASAR-Rep |
| --- | --- | --- | --- | --- |
| Chr19 | 0.933 | 0.969 | 0.982 | 0.980 |
| Chr20 | 0.914 | 0.956 | 0.975 | 0.978 |
| Chr21 | 0.908 | 0.917 | 0.976 | 0.972 |
| Chr22 | 0.927 | 0.969 | 0.983 | 0.983 |
| **NHEK** |  |  |  |  |
| Chr19 | 0.817 | 0.882 | 0.863 | 0.867 |
| Chr20 | 0.848 | 0.857 | 0.887 | 0.851 |
| Chr21 | 0.753 | 0.849 | 0.910 | 0.869 |
| Chr22 | 0.796 | 0.855 | 0.870 | 0.865 |
| **CH12LX** |  |  |  |  |
| Chr19 | 0.789 | 0.865 | 0.918 | 0.835 |
| Chr20 | 0.735 | 0.841 | 0.913 | 0.814 |
| Chr21 | 0.783 | 0.892 | 0.957 | 0.853 |
| Chr22 | 0.793 | 0.803 | 0.941 | 0.832 |

### 3.4 HiCSR effectively recovers TAD boundaries across varying sequence depths, cell types, and species

Finally, we evaluated HiCSR on a common Hi-C analysis task by exploring the effect of sequence depth and cell type on the model’s ability to recover contact domain boundaries. From the GM12878 cell type, we evaluated on chromosomes 19-22, from NHEK chromosomes 9, 13, 16 and 19, and from CH12LX chromosomes 8, 12, 16, and 18. We began by computing the normalized insulation scores (Crane *et al*., 2015) of the low resolution, HiCSR enhanced, and high resolution contact maps for each chromosome. We then compared the insulation scores of the low resolution and HiCSR enhanced data to the high resolution insulation scores using MSE.

We found that HiCSR enhanced data significantly reduces the error in the insulation score compared to low resolution data, and that the HiCSR framework is capable of recovering the insulation score values at a variety of sequence depths. In the GM12878 cell type, where the low resolution Hi-C data has a sequence depth of 407.80 M reads, a MSE of 0.012 was found for the low resolution insulation score, while the HiCSR enhanced insulation score achieves a MSE of 0.005. In the NHEK cell type (67.08 M reads), a MSE of 0.230 was found for the low resolution insulation score while the HiCSR enhanced score reaches a MSE of 0.103. Finally, for the CH12LX cell type (86.43 M reads), a MSE of 0.086 was found for the low resolution score, while the HiCSR enhanced data achieves a MSE of 0.040.

A sample of the contact maps, insulation scores, and resultant TAD boundaries (derived from the high resolution sample), for each cell type is illustrated in Fig. 7A. It can be seen that in all cases HiCSR is able to recover a sensible approximation of the insulation score. Even in cases where a lower sequence depth results in an extremely noisy signal, HiCSR enhanced data produced an insulation score which preserves the local minima used for TAD boundary annotation.

**Fig. 7.**
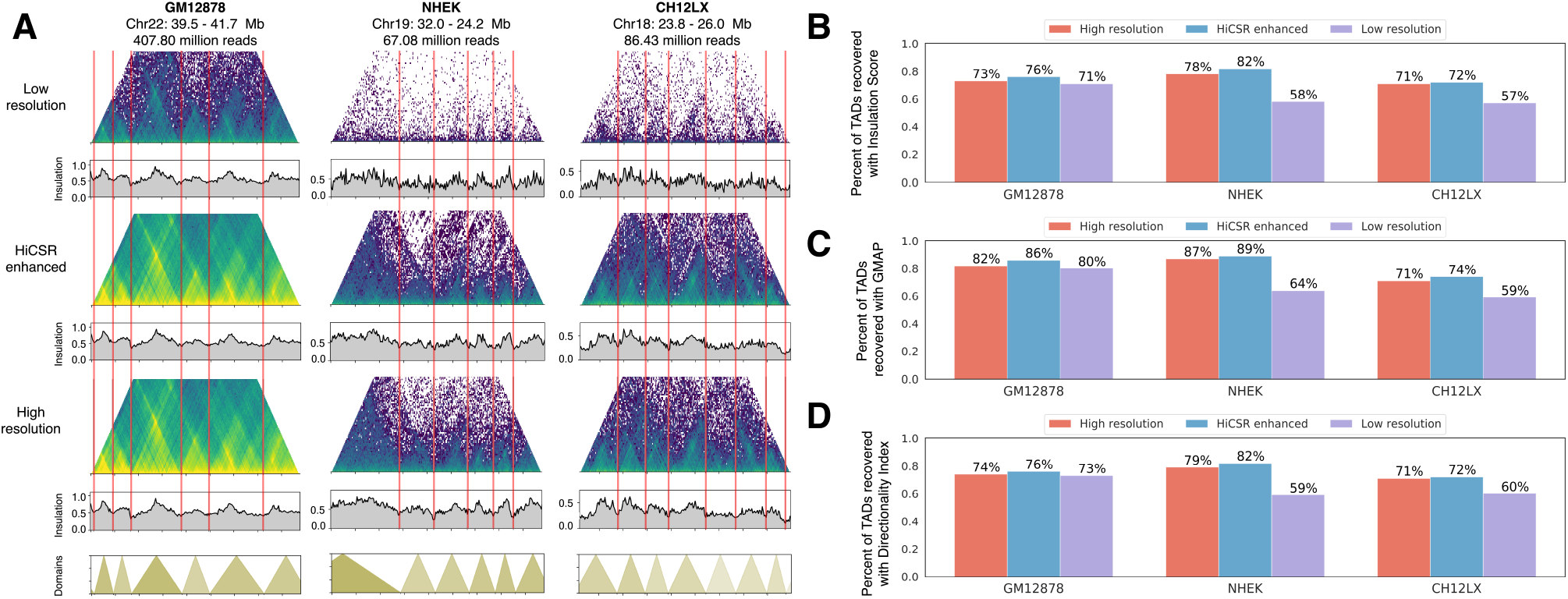
Evaluation of TAD boundaries recovered from HiCSR predicted Hi-C data across cell and species type. (A) Low resolution, HiCSR enhanced, and high resolution Hi-C data samples with corresponding insulation scores for the GM12878 and NHEK (human) cell types, as well as the CH12LX (mouse) cell type. The contact domain boundaries are shown with the boundary annotations highlighted in red for all three cell types. (B-D) Comparison of detected contact domain boundaries for each contact map relative to a curated set of known TAD boundary locations derived from the Insulation Score, Directionality Index, and GMAP methods.

We further validated HiCSR’s capability to recover TAD boundaries across sequence depth, cell type, and species by comparing TADs called from the insulation scores of the low resolution, HiCSR enhanced and high resolution data, to a known set of curated TAD boundaries from the TAD Knowledge Base (TADKB) (Liu T. *et al*., 2019) for each cell type. We consider over 1000 TAD boundaries from this knowledge base computed using the Insulation Score, Directionality Index (Dixon *et al*., 2012), and the Gaussian Mixture model And Proportion test (GMAP) (Yu *et al*., 2017) methods and determined the extent to which they could be recovered by HiCSR (Fig. 7 (B-D)). We deemed called contact boundaries to be correct if they fell within 0.16 Mb of a true contact boundary. We found that in all cases, HiCSR was able to call TAD boundaries more accurately than the original low resolution Hi-C data. In the GM12878 cell type, We also note that in both the NHEK and CH12LX cell types where the sequence depth is particularly low, HiCSR was able to recover a significant proportion of the true TAD boundaries, correctly locating over 20% more contact domain boundaries on average compared to the low resolution counterpart. Even within the GM12878 Hi-C data set with a much greater sequencing depth, we found that while the low resolution data was able to call TADs relatively better, it was still outperformed by the HiCSR enhanced insulation score.

## 4 Discussion

In this work, we showcase the HiCSR framework for enhancing low resolution Hi-C data. HiCSR is capable of producing highly accurate and visually convincing high resolution Hi-C contact maps from low resolution data. We believe that HiCSR will find use within the research community when analyzing low resolution Hi-C data obtained from either new experiments or online repositories. Our method leverages the strengths of all previously proposed deep learning methods and improves upon them by introducing a Hi-C specific feature reconstruction loss derived from a denoising autoencoder. To motivate the use of the denoising autoencoder for Hi-C representation learning, we first showed its capability in producing high fidelity reconstructions of Hi-C data. Furthermore, we demonstrated that the learned feature representations contain useful textures and interpretable Hi-C specific substructures including self-interacting loci, TADs, and chromatin loops.

Additionally, we believe that the denoising autoencoder developed to augment HiCSR’s objective function provides an interesting approach to task specific image based problems beyond super-resolution methods. In non-natural image based biological problems, it is not always clear which features of the image are important or useful. In these cases, the features learned by the denoising autoencoder provide an automated way to develop problem specific insights and loss functions which can be validated post-hoc by inspecting the model’s learned representations. We encourage future works to further explore these methods for unsupervised representation learning with both Hi-C data and biological data more broadly.

We argued that current evaluation methods of Hi-C enhancement (e.g. correlation methods) fall short as they do not consider the unique structures found within Hi-C data when evaluating the relative efficacy of Hi-C enhancement methods. We instead proposed the use of Hi-C specific measures of reproducibility and similarity which have been developed and endorsed by the Hi-C research community. We showed that HiCSR outperforms all previously proposed models achieving consistently high reproducibility scores measured with four separate metrics, indicating that HiCSR enhanced data is highly similar to the true high resolution Hi-C data from several reproducibility perspectives. Similarly, we found that HiCSR was capable of learning Hi-C super-resolution mappings which transferred well across cell types and again achieved high reproducibility scores. This indicates that there are shared properties underlying Hi-C contact maps across both cell types and species which are captured in HiCSR’s model parameters.

We also validated HiCSR’s performance in a common Hi-C analysis task, evaluating the model’s ability to recover TAD boundaries which are not detectable in low resolution data. We show that HiCSR is able to perform well at this task, providing robust results across a range of cell types, and between both human and mouse species. This indicates that HiCSR has captured the underlying structure which is shared in Hi-C data across varying cell types.

As Hi-C super-resolution methods are still relatively new, there are many avenues which may be explored to further improve the methodology. For example, one could investigate the use of ensemble methods. Training HiCSR on datasets from a variety of cell types, thereby increasing dataset size and variation, will likely improve cross-cell super-resolution predictions. Another possible improvement to our work is to provide side information to the model during training and when making predictions. For example, side information pertaining to the genomic distance of the current submatrix to be predicted may provide beneficial results, improving the model’s optimization behaviour. In addition to improving upon the methodology of HiCSR, work exploring the relative trade-offs between deep learning Hi-C enhancement methods (such as those discussed here) and recently proposed alternative methods such as HIFI (Cameron *et al*., 2020) is required to better understand the scenarios in which each technique is preferred.

Our contribution further improves upon the state-of-the-art in Hi-C super-resolution, providing a solution which produces highly realistic and accurate contact maps. We believe that continued refinements to the Hi-C enhancement process combined with more biologically relevant evaluations, as exemplified by this work, will accelerate its adoption by experimental biologists and contribute to our understanding of the 3D genome.

