## Supplementary Information for "HiCSR: a Hi-C super-resolution framework for producing highly realistic contact maps"

**Supplementary Table 1.** HiCSR framework network parameters. Additional parameters required to reproduce the generator ($G$), discriminator ($D$) and denoising autoencoder (DAE, $\phi$), including the number of input and output convolutional channels, kernel sizes, and stride.

| Layer | Layer type | Input channels | Number of kernels | Kernel size | Stride |
|---|---|---|---|---|---|
| **Generator,** $G$ | | | | | |
| 1 | Conv | 64 | 64 | $(3 \times 3)$ | 1 |
| 2 | Residual | 64 | 64 | $(3 \times 3)$ | 1 |
| ... | ... | ... | ... | ... | ... |
| 16 | Residual | 64 | 64 | $(3 \times 3)$ | 1 |
| 17 | Conv | 64 | 128 | $(3 \times 3)$ | 1 |
| 18 | Conv | 128 | 128 | $(3 \times 3)$ | 1 |
| 19 | Conv | 128 | 256 | $(3 \times 3)$ | 1 |
| 20 | Conv | 256 | 256 | $(3 \times 3)$ | 1 |
| 21 | Conv | 256 | 1 | $(3 \times 3)$ | 1 |
| **Discriminator,** $D$ | | | | | |
| 1 | Conv | 1 | 64 | $(4 \times 4)$ | 2 |
| 2 | Conv | 64 | 128 | $(4 \times 4)$ | 2 |
| 3 | Conv | 128 | 256 | $(4 \times 4)$ | 2 |
| 4 | Conv | 256 | 512 | $(4 \times 4)$ | 2 |
| 5 | Conv | 512 | 1 | $(1 \times 1)$ | 1 |
| **DAE,** $\phi$ | | | | | |
| 1 | Conv | 1 | 64 | $(3 \times 3)$ | 2 |
| 2 | Conv | 64 | 64 | $(3 \times 3)$ | 1 |
| 3 | Conv | 64 | 64 | $(3 \times 3)$ | 1 |
| 4 | Conv | 64 | 64 | $(3 \times 3)$ | 1 |
| 5 | Conv | 64 | 64 | $(3 \times 3)$ | 1 |
| 6 | Deconv | 64 | 64 | $(3 \times 3)$ | 1 |
| 7 | Deconv | 64 | 64 | $(3 \times 3)$ | 1 |
| 8 | Deconv | 64 | 64 | $(3 \times 3)$ | 1 |
| 9 | Deconv | 64 | 64 | $(3 \times 3)$ | 1 |
| 10 | Deconv | 64 | 1 | $(3 \times 3)$ | 2 |

**Supplementary Table 2.** Reproducibility scores from GenomeDISCO, HiC-Spector, HiCRep, and QuASAR-Rep for all comparison models in chromosomes 19-22, as well as X and Y of the GM12878 cell type. All reproducibility scores are computed between true high resolution and enhanced Hi-C data.

| **Chromosome 19** | GenomeDISCO | HiC-Spector | HiCRep | QuASAR-Rep |
|---|---|---|---|---|
| HiCPlus | 0.854 | 0.949 | 0.976 | 0.979 |
| HiCNN | 0.916 | 0.948 | 0.981 | 0.978 |
| hicGAN | 0.742 | 0.912 | 0.967 | 0.958 |
| DeepHiC | 0.716 | 0.569 | 0.980 | 0.955 |
| HiCSR (ours) | **0.933** | **0.969** | **0.982** | **0.980** |
| **Chromosome 20** | | | | |
| HiCPlus | 0.838 | 0.808 | 0.965 | 0.975 |
| HiCNN | 0.908 | 0.799 | 0.973 | 0.974 |
| hicGAN | 0.715 | 0.905 | 0.958 | 0.955 |
| DeepHiC | 0.731 | 0.891 | 0.974 | 0.956 |
| HiCSR (ours) | **0.914** | **0.956** | **0.975** | **0.978** |
| **Chromosome 21** | | | | |
| HiCPlus | 0.849 | 0.920 | 0.966 | 0.963 |
| HiCNN | 0.905 | **0.944** | 0.974 | 0.958 |
| hicGAN | 0.366 | 0.536 | 0.937 | 0.912 |
| DeepHiC | 0.712 | 0.544 | **0.979** | 0.950 |
| HiCSR (ours) | **0.908** | 0.917 | 0.976 | **0.972** |
| **Chromosome 22** | | | | |
| HiCPlus | 0.872 | 0.926 | 0.976 | 0.979 |
| HiCNN | 0.918 | 0.946 | 0.982 | 0.979 |
| hicGAN | 0.571 | 0.799 | 0.960 | 0.960 |
| DeepHiC | 0.656 | 0.815 | **0.983** | 0.959 |
| HiCSR (ours) | **0.927** | **0.969** | **0.983** | **0.983** |
| **Chromosome X** | | | | |
| HiCPlus | 0.853 | 0.856 | 0.895 | 0.974 |
| HiCNN | 0.911 | 0.773 | 0.918 | 0.973 |
| hicGAN | 0.598 | 0.788 | 0.955 | 0.947 |
| DeepHiC | 0.703 | 0.704 | **0.979** | 0.950 |
| HiCSR (ours) | **0.933** | **0.917** | 0.972 | **0.978** |
| **Chromosome Y** | | | | |
| HiCPlus | 0.919 | 0.889 | 0.960 | 0.870 |
| HiCNN | 0.912 | 0.887 | 0.931 | 0.963 |
| hicGAN | 0.776 | 0.899 | 0.948 | 0.966 |
| DeepHiC | 0.723 | 0.873 | **0.972** | 0.966 |
| HiCSR (ours) | **0.945** | **0.974** | 0.952 | **0.987** |

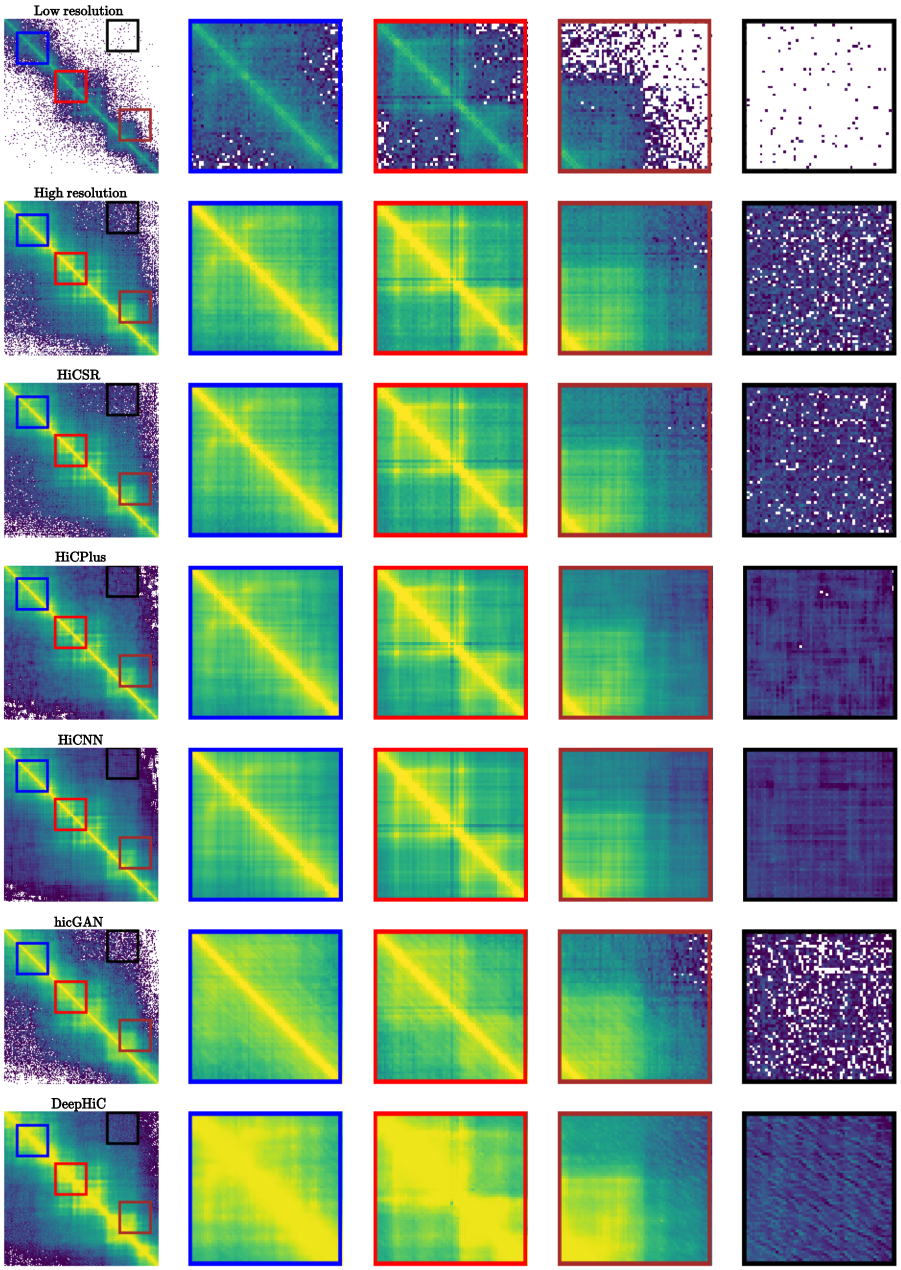


**Supplementary Fig. 1.** Visual samples of all Hi-C enhancement models compared, taken from chr22: 41.00 - 44.00 Mb in the GM12878 cell type. Blue, red, brown, and black boxes are close-ups taken from the left most image in each row. This example emphasizes the different enhancement biases and artifacts produced by each model.

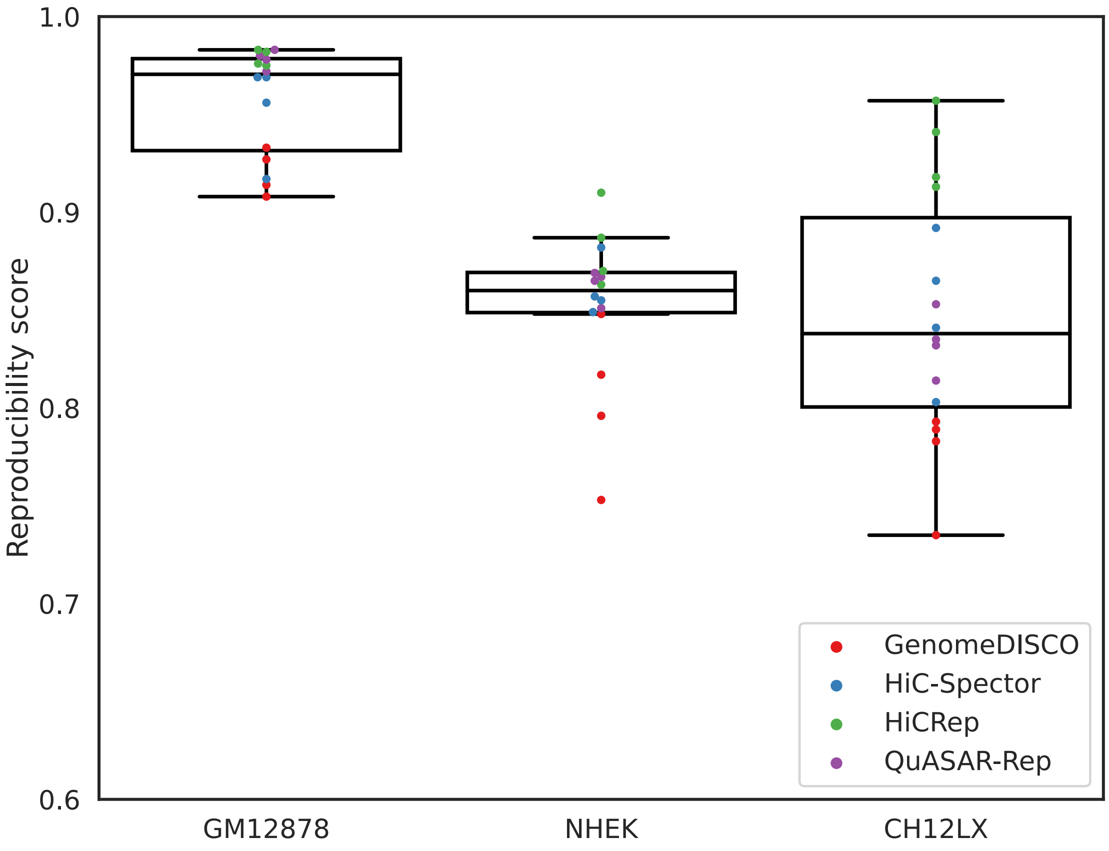


**Supplementary Fig. 2.** Comparison between HiCSR model performance on the GM12878 cell type when trained on datasets derived from the GM12878, NHEK, and CH12LX cell types. For each model, the four reproducibility metrics, GenomeDISCO, HiC-Spector, HiCRep, and QuASAR-Rep are computed.
